# The anti-tumor effect of trifluridine via induction of aberrant mitosis is unaffected by mutations modulating p53 activity

**DOI:** 10.1101/2023.08.30.555463

**Authors:** Takeshi Wakasa, Makoto Iimori, Kentaro Nonaka, Akihito Harada, Yasuyuki Ohkawa, Chie Kikutake, Mikita Suyama, Takashi Kobunai, Kazuaki Matsuoka, Kenta Tsunekuni, Yuki Kataoka, Hiroaki Ochiiwa, Kazutaka Miyadera, Takeshi Sagara, Eiji Oki, Shigehiro Ohdo, Yoshihiko Maehara, Hiroyuki Kitao

## Abstract

The fluorinated thymidine analogue trifluridine (FTD) is a chemotherapeutic drug commonly used to treat cancer; however, the mechanism by which FTD induces cytotoxicity is not fully understood. In addition, the effect of gain-of-function (GOF) missense mutations of the *TP53* gene (encoding p53), which promote cancer progression and chemotherapeutic drug resistance, on the chemotherapeutic efficacy of FTD is unclear. Here, we revealed the mechanisms by which FTD induced aberrant mitosis and contributed to cytotoxicity in both p53-null and p53-GOF missense mutant cells. In p53-null mutant cells, FTD induced DNA double-stranded breaks, single-stranded DNA accumulation, and the associated DNA damage repair responses during G2 phase. Nevertheless, FTD-induced DNA damage and the related responses were not sufficient to trigger strict G2/M checkpoint arrest. Thus, these features were carried over into mitosis, resulting in chromosome breaks and bridges, and subsequent cytokinesis failure. Improper mitotic exit eventually led to cell apoptosis, caused by the accumulation of extensive DNA damage and the presence of micronuclei encapsulated in the disrupted nuclear envelope. Upon FTD treatment, the behavior of the p53-GOF-missense-mutant, isogenic cell lines, generated by CRISPR/Cas9 genome editing, was similar to that of p53-null mutant cells. Thus, our data suggest that FTD treatment overrode the effect on gene expression induced by p53-GOF mutants and exerted its anti-tumor activity in a manner that was independent of p53 function.

## INTRODUCTION

Nucleoside analogues are commonly used in the treatment of cancer (1). These anti-cancer drugs act as anti-metabolites, compete with physiological nucleosides, and induce various intracellular responses, including p53 activation, to exert cytotoxic effects (2). Trifluridine (FTD), which belongs to the fluoropyrimidine class of drugs and is a component of the clinically approved chemotherapeutic drug called trifluridine/tipiracil (FTD/TPI) (3-5), is efficiently internalized by tumor cells, where it is rapidly phosphorylated by nucleoside kinases, including thymidine kinase 1 (6), to triphosphate. The triphosphate is then incorporated into DNA strands during DNA replication. Incorporation of FTD into DNA induces retarded replication fork progression (termed DNA replication stress) and the associated cellular response to stress (7,8). Although FTD-induced, p53-status-independent growth suppression has been reported (7,9), its effect was typically determined by different mechanisms in a p53-status-dependent manner. Under conditions of FTD-induced DNA replication stress, tumor cells expressing wild-type p53 exhibit p53–p21 pathway activation, cyclin B1 degradation during G2 phase, mitotic skipping, and subsequently cellular senescence. However, we have previously shown that mitotic entry, aberrant chromosome segregation, and apoptosis occurred in isogenic p53-null mutant cells (7). A comprehensive analysis of variations in the *TP53* gene, which encodes the tumor suppressor protein p53, revealed that *TP53* missense mutations are the most prevalent in human cancer (10). Furthermore, 95% of *TP53* missense mutations occur in the core (DNA-binding) domain of p53. Six p53 residues, namely, R175, G245, R248, R249, R273, and R282, have been identified as mutation hotspots (11). Although the expression of full-length p53 is unaffected by these mutations, they abolish its tumor suppressor function (12). Recent studies have revealed that these p53 variants acquire neomorphic oncogenic activities, which results in abnormal cell growth (13), perturbation of metabolic pathways (12,14), and tumor metastasis (15,16). Furthermore, many cancer studies have shown that such *TP53*-targeting, gain-of-function (GOF) mutations contribute to resistance of cells to radiation and chemotherapeutic drugs such as cisplatin, doxorubicin, etoposide, and 5-FU (17-20). However, the effect of *TP53*-GOF mutations on the chemotherapeutic efficacy of FTD remains to be elucidated. In addition, the mechanisms underlying FTD-induced DNA replication stress in p53-deficient cells, which results in abnormal mitotic progression and subsequent cell death, are unclear.

In this study, we reported that FTD-induced DNA replication stress contributed to cytotoxicity via a mechanism that depended on the loss of p53 function. We further used a panel (constructed by CRISPR/Cas9 genome editing), comprising p53-wild-type, p53-null, and p53-GOF mutant cells, to demonstrate that the anti-tumor effect of FTD occurred independently of p53 status.

## MATERIALS AND METHODS

### Cell culture and reagents

HCT116 (ECACC) cells were cultured in DMEM (Thermo Fisher). LS174T (ATCC) and WiDr (ATCC) cells were cultured in EMEM (Thermo Fisher). LS1034 (ATCC) and COLO 320DM (ATCC) cells were cultured in RPMI1640 (Thermo Fisher). All media were supplemented with 10% fetal bovine serum. All cell lines were grown in a 5% CO_2_ atmosphere at 37°C. All cell lines were authenticated by short tandem repeat (STR) analysis and confirmed negative for *Mycoplasma* contamination. The final concentration of each reagent was as follow: trifluorothymidine (FTD) : 3 µM (T2511; Tokyo Chemical Industry), 5-fluorouracil (5-FU) : 3 µM (F6627; Sigma), cisplatin (CDDP) : 5 µM (D3371; Tokyo Chemical Industry), aphidicolin : 0.5µM (A0781; Sigma); neocarzinostatin (NCS) : 20 nM (N9162; Sigma); RO-3306: 9 µM (SML-0569; Sigma).

### Generation of p53 missense mutant cells

To knock-in *TP53*-missense mutation using the CRISPR/Cas9 genome editing system, guide RNA (gRNA) sequences were designed using the online software CRISPRdirect

(21). The sense and antisense oligonucleotides were annealed and cloned into the *Bbs*I site of pX330A-1x2 (Addgene; plasmid #58766), which was a gift from Feng Zhang. To construct donor vectors, left- and right-arm DNA fragments were amplified using template DNA (genomic DNA from HCT116 cells), KOD FX DNA polymerase (TOYOBO) and primers containing approximately 500 base pairs sequence homologous to the target locus and then cloned into the *EcoR*V site of pBluescript SK+ using an In-Fusion HD Cloning Kit (Takara Bio). *TP53*-missense mutations and protospacer adjacent motif (PAM) sequence mutation were introduced using the QuikChange Lightning site-directed mutagenesis kit (Agilent Technologies). A puromycin or neomycin resistance cassette was cloned into the *BamH*I site which was placed between left- and right-arm of the amplified and cloned DNA fragments. The sequences of the gRNAs and primers were provided in supplementary table.

To establish the *TP53*-missense mutation knock-in cell lines, as shown in supplementary fig. 2, HCT116 cells were co-transfected with the pX330 plasmid containing the single gRNA target sequence and two donor DNA plasmids harboring puromycin or neomycin resistance cassette using a 4D Nucleofector (Lonza). After double selection with puromycin and G418 (geneticin) (Thermo Fisher), each clone was screened by genomic PCR and sequencing. The puromycin and neomycin resistance cassettes integrated into the *TP53* loci were removed by the Cre-*loxP* recombination system. Cre recombinase proteins were delivered into *TP53*-missense mutation knock-in cells using Cre Recombinase Gesicles (Takara Bio).

### Generation of fluorescent ubiquitination-based cell cycle indicator (Fucci)- expressing cells

A PIP-FUCCI coded DNA fragment was PCR amplified with template DNA (pENTR-PIP-FUCCI; Addgene; plasmid #118621), which was a gift from Jean Cook, KOD FX DNA polymerase (TOYOBO) and primers (supplementary table 1) and then cloned into the *BamH*I site of pLVSIN-EF1 α Hyg vector (Takara Bio) using an In-Fusion HD Cloning Kit (Takara Bio). To produce lentiviral stocks, lenti-X 293T Cells (Takara Bio) were co-transfected with Lentiviral High Titer Packaging Mix (Takara Bio) and the lentiviral plasmid (pLVSIN-PIP-FUCCI or pBOB-EF1-FastFUCCI (Addgene; plasmid # 86849), which was a gift from Kevin Brindle & Duncan Jodrell), using a TransIT-Lenti Transfection Reagent (Takara Bio). Viral solutions were collected 48 h post-transfection, filtered through a Millex-HV 0.45 µm low protein binding PVDF membrane (Millipore), and concentrated 100-fold using Lenti-X Concentrator (Takara Bio) following the manufacturer’s protocol. HCT116 cell lines were infected to generate a polyclonal population expressing Fucci SA or PIP-Fucci and sorted proliferating cells using cell sorter SH800 (SONY).

### Quantitative reverse-transcription PCR (qRT-PCR)

Total RNA was extracted from each cell line using a RNeasy Mini Kit (Qiagen). cDNA was synthesized with a High-Capacity cDNA Reverse Transcription Kit (Applied Biosystems) using the primers provided in Supplementary Table 1. mRNA expression was normalized against that of *β-actin*. Quantitative RT-PCR was performed with a THUNDERBIRD SYBR qPCR Mix (TOYOBO). Fluorescence signals were detected by a LightCycler 480 system (Roche Diagnostics).

### RNA-seq and gene expression analysis

For RNA-seq analysis of HCT116 wild-type or mutant cells, six replicates were performed for each two independent clones. The cells were cultured to mid-log phase and collected for extracting total RNA. RNA-seq library was prepared with CEL-seq2 technique as described previously (22). The CEL-seq2 library was sequenced on an Illumina HiSeq 1500 system (Illumina) with the following cycles: 15 cycles for read 1 and 45 cycles for read 2 (23). Paired-end reads were generated using this platform. Sequence reads in FASTQ format was applied to FastQC (v0.11.9) to assess sequence quality. For adapter trimming, Trim Galore! (v0.6.0) was used. The sequence reads were mapped to human reference genome GRCh38/hg38 from Ensembl (release 84) using HISAT2 (version 2.1.0). The mapped reads were counted for genes using featureCounts (v.1.6.4). The mapped data were converted to counts per million (CPM) and log2-transformed. The expression profiles were compared using the unpaired Student *t*-test with the Benjamini and Hochberg procedure for controlling the false discovery rate (FDR) (24) and a fold change value. Genes with an adjusted FDR <0.005 and absolute values of log FC > 1 were considered to be differentially expressed genes. Two-dimensional hierarchical clustering was then applied to the log-transformed data with centroid-linkage clustering with the Pearson correlation-based distance as the similarity metric for the discriminating genes identified as differentially expressed according to each p53 status. Variations in the multigene expression among each p53 status were also compared by a principal component analysis (PCA). The first 2 principal components were fitted to the cell’s molecular profile data and plotted into a 2-dimensional space. The cumulative proportion of the variance captured by each principal component axis was also calculated. These data analysis was performed in the R computing environment and GeneSpring GX software version 13.0 (Agilent Technologies).

### Western blotting

Cells were harvested and lysed in RIPA buffer (50 mM Tris-HCl pH 8.0, 150 mM NaCl, 0.1% SDS, 1% NP-40, 0.5% deoxycholate, 1 mM PMSF and a protease and phosphatase inhibitor cocktail (Nacalai Tesque)) for 30 min on ice, and then sonicated. Cell extracts were clarified by centrifugation. Cell lysates were boiled in Laemmli sample buffer (Bio-Rad). Cell lysates were separated by SDS-PAGE with TGX precast gels (Bio-Rad) and transferred to membranes using the Trans-Blot Turbo Transfer System (Bio-Rad). Western blotting was performed using the primary and secondary antibodies provided in Supplementary Table 2. Detection was performed using the Chemi-Lumi One (Nacalai Tesque) with a LAS 4000 mini (Cytiva).

### Co-immunoprecipitation

The ETS2 gene was PCR amplified from HCT116 cDNA using KOD FX DNA polymerase (TOYOBO) and primers (supplementary table 1), cloned into pENTR/D-TOPO (Thermo Fisher), and then transferred into the pcDNA3.1 vector (Gateway Technology, Thermo Fisher) harboring 3FLAG for tagging at N-terminus. HCT116 wild-type or p53 missense mutant cells were transfected with the plasmids using X-treme GENE HP (Merck) according to the manufacturer’s instructions. After 24 h, cells were washed with cold PBS, lysed in IP buffer (20mM Tris-HCl, pH 8.0, 137mM NaCl, 1mM MgCl_2_, 1mM CaCl_2_, 1% NP-40, 10% glycerol, 1 mM PMSF, a protease and phosphatase inhibitor cocktail (Nacalai Tesque), and 12.5U ml^-1^ benzonase (Millipore)), and sonicated. After centrifugation, the supernatants were collected and incubated with 50 μl of anti-DDDDK-tag mAb-Magnetic Beads or Mouse IgG2a Magnetic Beads (MBL) at 4°C for 2 h. Beads were pelleted by a magnetic rack, washed 5 times in 1 mL IP buffer, resuspended to 30μL Laemmli sample buffer (Bio-Rad), and boiled. Immunoprecipitated proteins were analyzed by immunoblotting as already described.

### Immunofluorescence

Cells were grown on collagen-I-coated coverslips (Iwaki). For staining of Fucci SA and PIP-Fucci, cells were fixed in 4% paraformaldehyde at 37°C for 10 min and then in 90% methanol at room temperature for 5 min. For staining of other proteins, cells were fixed in 4% paraformaldehyde at 37°C for 15 min. Thereafter, cells were rinsed with phosphate-buffered saline (PBS) containing 10 mM glycine, permeabilized in PBS containing 0.1% Triton X-100 for 5 min, blocked in PBS containing 2% bovine serum albumin (BSA) for 30 min at room temperature and incubated with the primary and secondary antibodies provided in Supplementary Table 2. Coverslips were washed with PBS containing 4,6-diamidino-2-phenylindole (DAPI) for 5 min, washed with PBS and mounted with Prolong Glass (Thermo Fisher).

### Chromosome spreads

Cells were treated with FTD for 48h, and then with 0.1 μg/mL colcemid (Karyomax; Thermo Fisher) for 12 h. Chromosome spreading was performed as described previously (25).

### Spheroid culture

The cells were seeded into a 96-well spheroid culture plate (EZ-BindShut II; IWAKI) with the number of 300 cells per well, incubated for 24 h at 37 °C, and then treated with 9 μM FTD. After 3, 6 and 9 days of FTD treatment, the viability of spheroid cells was assessed with Cell-titer GLO 3D cell viability assay (Promega) and TriStar LB 941 plate reader (Berthold Technologies).

### Image acquisition and analysis

For fixed-cell experiments, fluorescence image acquisition was performed using a Nikon A1R confocal imaging system controlled by the Nikon NIS Elements software (Nikon). The objective lens was an oil immersion Plan-Apo 100× numerical aperture (NA) 1.40 lens (Nikon). Images were acquired as Z-stacks at 0.2-mm intervals and deconvoluted. Maximum-intensity projections were generated using the NIS Elements software (Nikon). For live-cell imaging, cells stably expressing Fucci or histone H2B-GFP were grown in a glass bottom chamber (Matsunami) that had been coated with collagen-I (Nippi) in 5 mM acetic acid at room temperature for 30 min before use. Cells were maintained at 37℃ in the presence of 5% CO_2_ in a stage-top incubator and PureBox SHIRAITO (Tokai Hit) in the following medium throughout the entire duration of live-cell imaging: FluoroBrite DMEM (Thermo Fisher) supplemented with 10% FBS and GlutaMAX (Thermo Fisher). Images were acquired every 10 min using a Plan-Apo 20× NA 0.75 lens (Nikon) on an inverted fluorescence microscope Eclipse Ti-E (Nikon) equipped with a DS-Qi2 camera (Nikon). For chromosome spreading experiments, an oil immersion Plan-Apo 100× numerical aperture (NA) 1.40 lens (Nikon) on a BZ-X800 fluorescence microscope (KEYENCE) was used for image acquisition. For spheroid observation, bright-field images were acquired with a CFI achromat 10×F NA 0.25 lens (Nikon) on an inverted microscope Eclipse TS100 (Nikon) equipped with a DS-Fi2 camera (Nikon).

### Xenograft

Male nude (BALB/cAJcl-nu/nu) mice were purchased from CLEA Japan, and were housed under specific pathogen-free conditions, at 23 ± 3°C and 50 ± 20% humidity on a 12-h light–dark cycle, with normal diet (CE-2; CLEA Japan) and water provided ad libitum. The mice were quarantined for one week and then subcutaneously implanted with 2x10^7^ cells of HCT116, HCT116 p53-KO, or HCT116 p53-R175H cells, or with 5x10^7^ cells of HCT116 p53-R248Q cells. The mice were randomized on day 0 based on approximately 200 mm^3^ of tumor volume and each treatment group consisted of six mice. FTD/TPI was prepared by mixing FTD and TPI at a molar ratio of 1:0.5 in 0.5% HPMC solution at 10 mL/kg. FTD/TPI (FTD: 200 mg/kg/day) was administered orally once daily from day 1 to 5 and day 8 to 12. Tumor volume was calculated as 1/2 × (length × width × width). All the animal studies were performed according to the guidelines and with the approval of the institutional Animal Care and Use Committee of Taiho Pharmaceutical Co., Ltd. Ethical approval (10 Oct 2019) was obtained prior to conducting the animal experiments.

### Data availability

RNA sequencing data for Fig. 3D and E have been deposited in the DDBJ Sequence Read Archive under accession number DRA014321.

## RESULTS

### FTD induces errors in mitotic progression and subsequent aberrant nuclear morphology in *TP53*-knockout cells

We first assessed the cell cycle status of FTD-treated *TP53*-knockout cells in a *TP53*-knockout HCT116 cell line (hereafter referred to as p53-KO cells), which was previously generated using the CRISPR/Cas9 genome editing system (7,26). To exclude the effect of different durations of FTD treatment in S phase, we synchronized the cells by inducing cell cycle arrest using thymidine and RO-3306 (a CDK1 inhibitor). We added FTD into cell culture media 5 h after release from RO-3306, when most cells were in G1 or early S phase (Fig. 1A). The accumulation of cells containing 4N DNA was measured 24–72 h after FTD administration. By contrast, vehicle treated cells containing 4N DNA were converted to 2N-DNA-containing ones within 15 h of vehicle administration. In a previous study, we showed that FTD induced extensive chromosome bridge formation in *TP53*-knockout cells following mitotic entry (7). Therefore, we speculated that the FTD-induced accumulation of 4N-DNA-containing cells might not only signal sustained G2 phase but also post-mitotic G1 phase. To investigate this possibility, we compared the mitotic progression of FTD-treated and untreated cells expressing histone H2B-GFP using live-cell imaging. FTD-treated cells were differentiated from control cells by exhibiting a longer duration between mitotic entry and anaphase onset, as well as abnormal mitotic exit (Fig. 1B and Supplementary movie). Of note, different types of defects were observed at late mitosis in the FTD-treated cells, such as unequal sister chromatid segregation, followed by abnormal anaphase with chromosome bridges (Fig. 1B, middle panel), and subsequent cytokinesis failure (Fig. 1B, lower panel). To confirm this phenomenon in detail, we used the fluorescent ubiquitination-based cell cycle indicator (Fucci SA) system to establish HCT-116 cells, which could be used to visualize cell cycle status (27). We used monomeric Kusabira Orange (mKO2)-fused human Cdt1 (hCdt1) (amino acid residues 30–120), which accumulates in Fucci-expressing cells during G1 phase, and found that most of the FTD-treated mKO2+ Fucci-expressing cells contained multi- or micronuclei, which were rarely observed in the control cells. Furthermore, about half of the FTD-treated cells containing multi- or micronuclei had two or more centrosomes in G1 phase (Fig. 1C, D). Taken together, our results suggest that FTD treatment induced chromosome mis-segregation or cytokinesis failure via chromosome bridge formation during mitosis, which perturbed nuclear formation and resulted in the generation of multi-or micronuclei.

**Figure 1.**
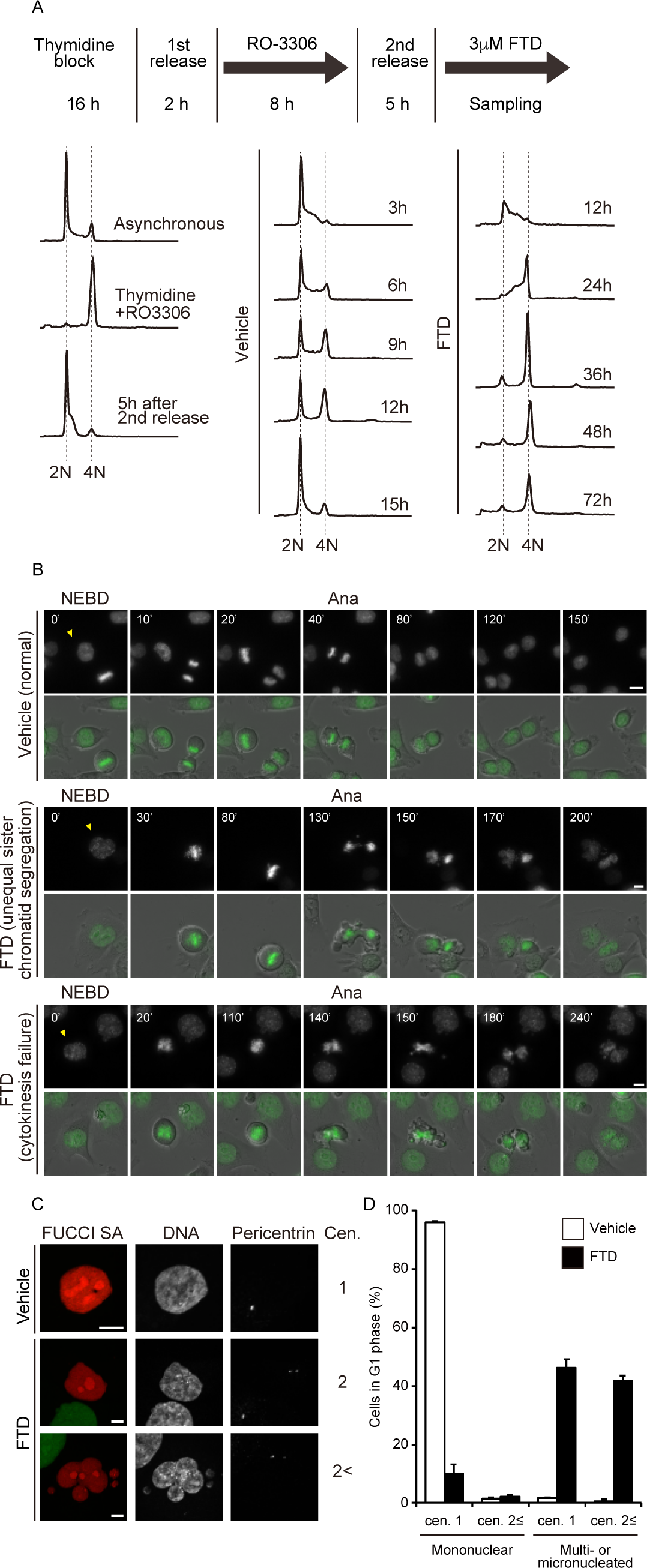
FTD promotes improper mitotic progression in p53 KO cells. A, Flow cytometry cell cycle analysis using propidium iodide (PI) DNA staining and its experimental scheme. Synchronous HCT116 p53 KO cells were treated with FTD or not and then collected at indicated time points after 2nd relase. B, Selected frames from live-cell imaging of representative HCT116 p53 KO cells expressing histone H2B-GFP treated with FTD. Time (minutes) after nuclear envelope breakdown (NEBD) is shown on the images. Arrowheads highlight representative cell images. Ana, onset of anaphase. Scale bar, 10 µm. C and D, HCT116 p53 KO cells expressing Fucci SA were treated with FTD, fixed and stained with the indicated antibodies. Representative images of G1 cells are shown in C. Quantification of cells in G1 phase with multiple centrosomes and aberrant nuclei formation is shown in D. Data are means± s.d. from three independent experiments (≥ 250 cells per experiment). Scale bar, 10 µm.

### FTD-induced DNA damage is maintained between G2 phase and the following G1 phase and causes chromosome breaks and aberrant morphology during mitosis

Although FTD induced aberrant mitosis and subsequent apoptosis in p53-KO cells (Fig. 1 and (7)), the mechanisms underlying the induction of chromosome segregation errors and post-mitotic cell death remained unclear. We previously reported that FTD treatment in S phase induced the accumulation of persistent DNA lesions, including single-stranded (ss)DNA (7). To further evaluate the extent of DNA damage and its repair response during G2 phase and mitosis, we generated HCT116 PIP-Fucci cells, which can be used to distinguish between S and G2 phase via the detection of phase transition (28). After treatment with FTD, the Cdt1 (amino acid residues 1–17)- mVenus/ mCherry-Geminin (amino acid residues 1–110)-double-positive HCT116 p53-KO cells (in G2 phase; yellow color) formed γH2AX and RPA32 nuclear foci, indicating the presence of DNA double-stranded breaks (DSBs) and ssDNA accumulation. Consistent with this observation, the activation of DNA non-homologous end joining (NHEJ) and homologous recombination (HR) repair pathways in G2 phase were markedly induced in these cells, as evidence by the formation of 53BP1 and RAD51 foci (Fig. 2A). Furthermore, the FTD-induced ssDNA accumulation, DSBs and the associated repair responses were relayed from interphase into mitosis, resulting in the localization of RPA32 foci and the co-localization of γH2AX/MDC1 foci on the condensed chromosomes (Fig. 2B and Supplementary Fig. 1A).

**Figure 2.**
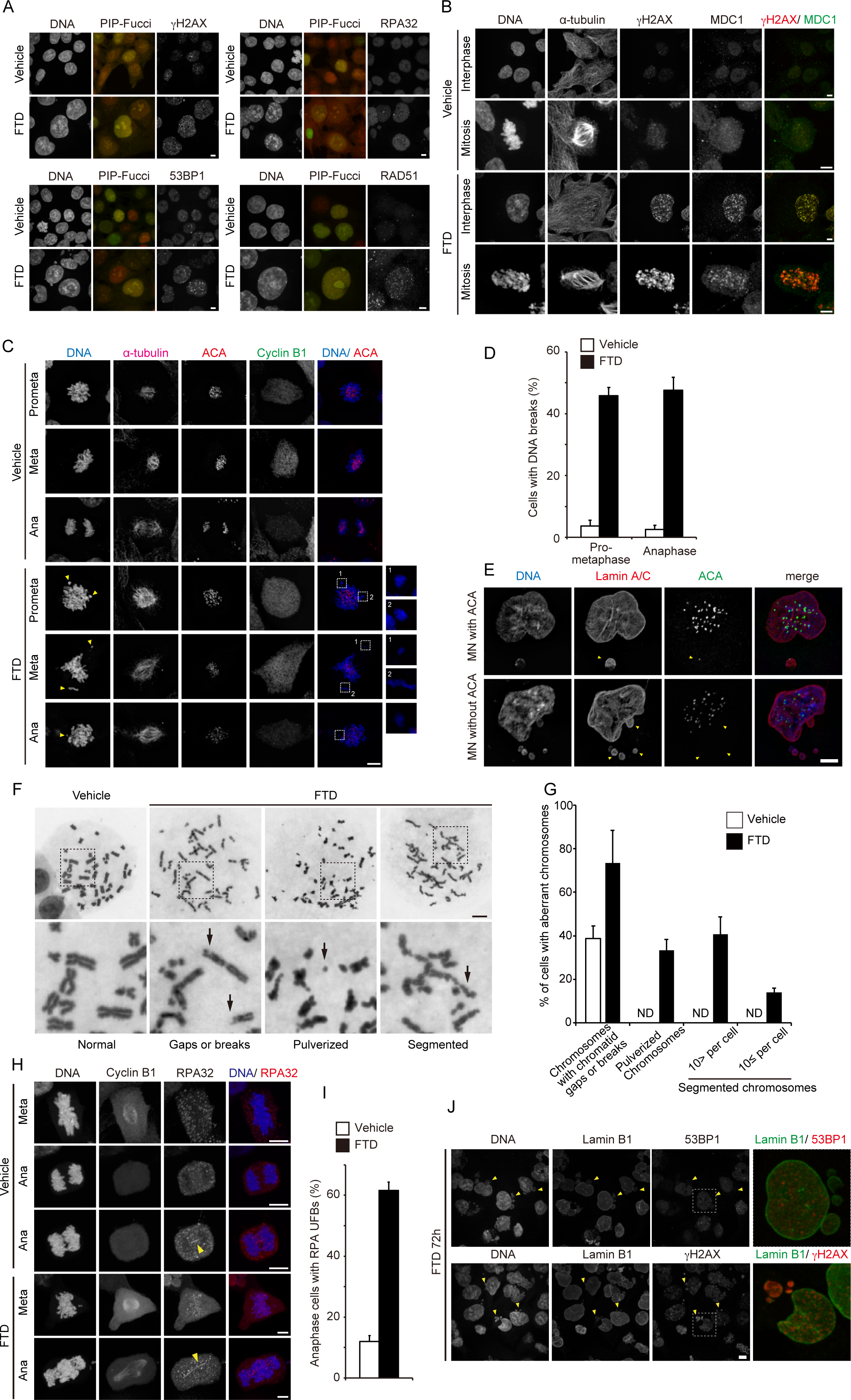
Effect of FTD treatment on mitotic progression in p53 KO cells. A, Representative images of cells in G2 phase. HCT116 p53 KO cells expressing PIP-Fucci were treated with FTD, fixed and stained with the indicated antibodies. Scale bar, 10 µm. B, Immunofluorescence images showing the localization of gH2AX/ MDC1 in FTD-treated mitotic cells. Cells were fixed and co-stained with the indicated antibodies. Scale bar, 10 µm. C and D, Representative images of broken chromosome arms without centromere signals during mitosis. Cells were treated with FTD, fixed and co-stained with the indicated antibodies. Arrowheads highlight broken chromosome arms. Quantification of broken chromosome arms is shown in D. Data are means± s.d. from three independent experiments (≥ 250 cells per experiment). E, FTD-treated post-mitotic cells. Cells were fixed and co-stained with the indicated antibodies. Arrowheads highlight micronuclei. Scale bar, 10 µm. F and G, Giemsa-stained chromosome spreads of FTD-treated cells. Representative images of normal or abnormal chromosomes are shown in F. Arrows highlight phenotypes of abnormal chromosome. The upper panels show whole chromosome spreads whereas the lower panels show the indicated sections at higher magnification. Quantification of abnormal chromosomes is shown in G. H and I, FTD-treated cells at anaphase with RPA32-positive UFBs were determined. Quantification of proportion of cells exhibiting UFBs is shown in I. Arrowheads highlight UFBs. Scale bar, 10 µm. J, FTD-treated post-mitotic cells. Cells were fixed and co-stained with the indicated antibodies. Arrowheads highlight micronuclei. Enlarged images show the indicated sections at higher magnification. Scale bar, 10 µm.

As the formation of γH2AX/MDC1 foci was highly prevalent in FTD-treated mitotic cells, we speculated that chromosome structure might be compromised during mitotic progression. Indeed, about half of the FTD-treated mitotic cells had broken chromosome arms, as evidenced by DNA condensation in the absence of centromere signals (Fig. 2C, D). These broken chromosome arms appeared as micronuclei lacking a centromere in the post-mitotic phase (Fig. 2E).

To confirm the presence of and characterize these structural chromosome abnormalities, we performed chromosome spreads of FTD-treated cells and found that a large number of cells had chromosome gaps, breaks, and pulverization. Of note, a high percentage of cells exhibited severe chromosomal abnormalities, such chromosomal elongation and segmentation (Fig. 2F, G). These phenomena looked very similar to the persistent mitotic HR intermediates, in which unresolved Holliday junctions arise from resolvase deficiency (29); these were also reported as a cause of HR ultra-fine DNA bridges (UFBs) (29). Indeed, FTD-treated cells displayed not only chromosome bridges (which can be visualized with DAPI staining) but also RPA-decorated UFBs at anaphase, which were rarely observed in the vehicle treated cells (Fig. 2H, I). Mitotic DNA synthesis, which is a mechanism used in the repair of under-replicated DNA by MUS81 and POLD3 (30), was rarely observed in FTD-treated mitotic cells, compared with aphidicolin-treated cells, even though FTD induced the accumulation of under-replicated regions and ssDNA (Supplementary Fig. 1B and (7)). Next, we examined how FTD-induced DNA damage, observed at G2 phase and mitosis, affected cell fate at the post-mitotic G1 phase. As indicated in Figure 1D, about 90% of the cells treated with FTD were multi- or micronucleated. Hatch *et al.* reported that mis-segregated mitotic chromosomes induce nuclear envelope rupture in the micronuclei, as a result of defects in nuclear lamina assembly. Furthermore, such nuclear envelope disruption markedly impairs nuclear functions (e.g., DNA damage repair mechanisms) and induces massive accumulation of damaged DNA (31). We found that FTD-induced micronuclei exhibited discontinuous lamin B1 localization at the micronuclear rim (Fig. 2J; arrow heads). Although 53BP1 foci (a DNA damage response factor) were observed in the primary nuclei of these cells, their foci were not detected in the disrupted micronuclei characterized by lamin gaps (Fig. 2J upper).

However, γH2AX accumulation in the form of a single large focus was seen in the disrupted micronuclei, suggestive of extensive DNA damage (Fig. 2J, lower panel). Taken together, our results suggest that FTD treatment not only triggered major DNA damage in G2 phase, but also perturbed mitotic progression via chromosome bridge formation. This may have led to HR intermediate generation, resulting in cumulative DNA damage and apoptosis induction in the next G1 phase.

### The effect of *TP53*-knock-in, GOF point mutations on genome-wide gene expression

We next investigated the effect of differential p53 status on the responsiveness of cells to anti-cancer drugs, especially FTD. However, comparison of p53 status among different tumor cell lines cannot rule out the effects of genetic variations other than p53 status. To overcome such limitations, we first generated *TP53*-mutant, isogenic cell lines by genome-editing HCT116 cells, which have wild-type p53 and near-diploid karyotypes, using the CRISPR/Cas9 system. In addition to the HCT116 p53-KO cells, we generated two classes of homozygous *TP53* GOF mutation knock-in HCT116 cell lines: p53-R175H and p53-R248Q (Supplementary Fig. 2). These mutations were classified into structural (R175H) and DNA-contacting (R248Q) mutations, which are the most frequently encountered p53-targeting mutations in human cancer (11,32).

We then assessed the properties of the cell lines harboring *TP53* missense mutations. The basal mRNA expression levels of the p53-targeted genes, *MDM2* and *CDKN1A* (encoding p21) (33), were significantly decreased in both the p53-R175H and p53-R248Q cells, as well as in the p53-KO cells (Fig. 3A). We next evaluated the response of the p53–p21 signaling pathway to DNA-damaging agents, including the nucleotide analogs FTD and 5-FU, the DNA-crosslinking agent cisplatin, the DNA replication inhibitor aphidicolin, and the DNA-breaking agent neocarzinostatin. Consistent with the results of basal *CDKN1A* expression, all the above agents failed to induce p21 accumulation, even though the p53 *per se* (in the p53-R175H, p53-R248Q, and p53 wild-type cells) was sensitive to all these DNA-damaging agents (Fig. 3B). These data suggest that the p53 missense mutant proteins exhibited a loss-of-function of wild-type p53 transcriptional activity in the p53-R175H and p53-R248Q mutant knock-in HCT116 cells.

**Figure 3.**
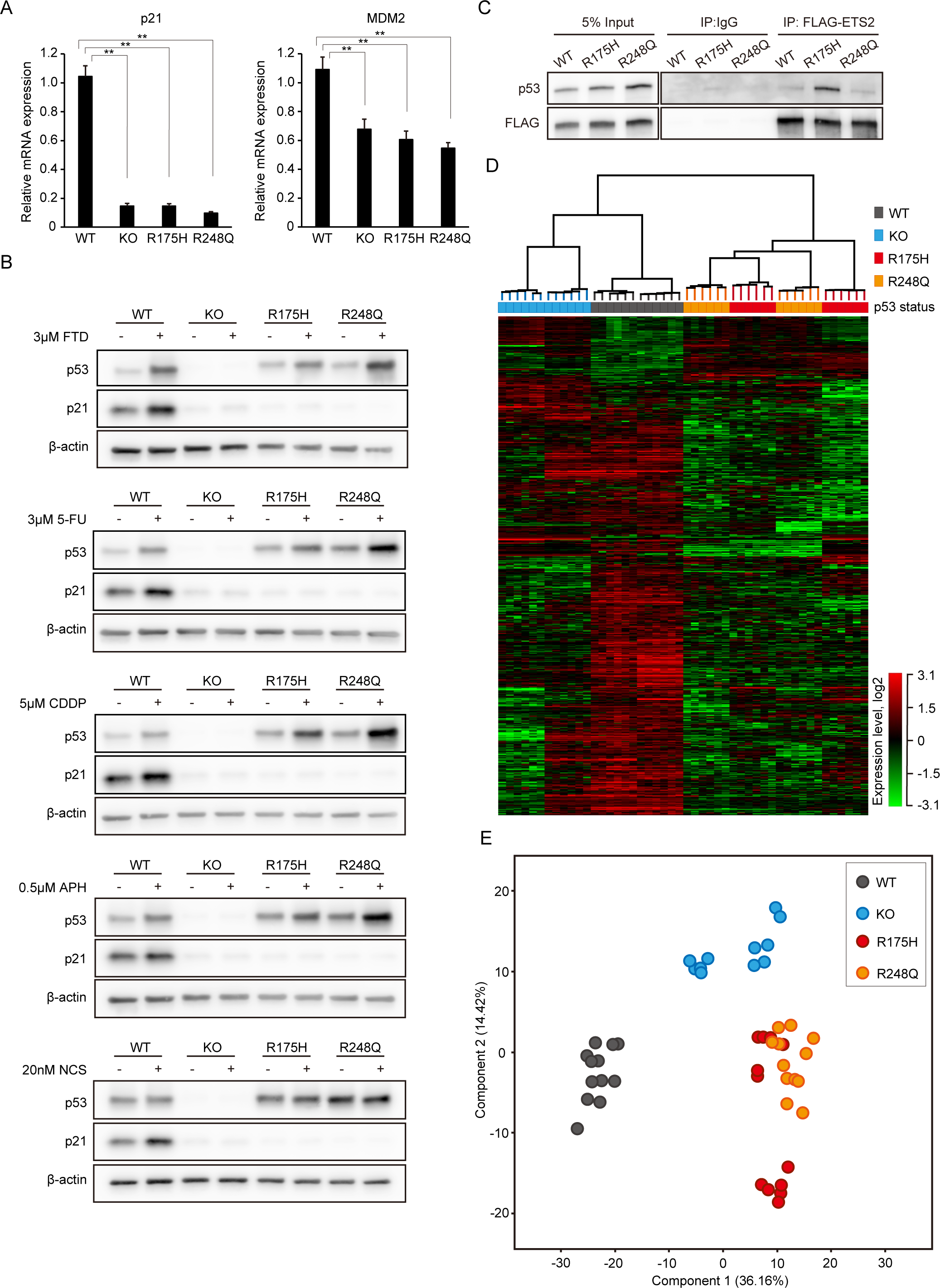
Generation of p53 GOF mutation knock-in cells. A, The basal mRNA expression levels of MDM2 and p21. Data are means± s.d. from three independent experiments. **P < 0.01. B, HCT116 cell lines were treated with the indicated drugs for 24 hrs. Western blot analysis was carried out using antibodies against the indicated proteins. C, Co-immunoprecipitation of HCT116 wild-type and GOF mutant cell-expressed Flag-ETS2 with endogeneous p53. Western blot analysis was carried out using antibodies against p53 proteins and FLAG tag. D and E, RNA-seq and gene expression analysis. For this analysis of HCT116 wild-type or mutant cells, six replicates were performed for each two independent clones. Two-way hierarchical clustering analysis and Principal components analysis among HCT116 cell lines harboring each indicated p53 status is shown in D and E, respectively. See materials and methods section for details.

Next, we assessed the function of the GOF mutant p53 in the p53-R175H and p53-R248Q cell lines. It was reported that p53 GOF missense mutations mapping to the DNA-binding domain, such as R175H or R248Q, cause p53 to associate with non-canonical transcription factors and trigger gene expression (12,18,34). For instance, ETS2, a E26 transformation-specific (ETS) family member, is known to associate with p53 missense mutants (18,34). Consistent with these reports, the results of our co-immunoprecipitation analysis demonstrated that ETS2 interacted with p53-R175H but not with wild-type p53 (Fig. 3C). Of note, ETS2 has also been shown to interact with p53-R248Q, albeit to a lesser extent than with p53-R175H or p53-R248W (18,34). Indeed, our co-immunoprecipitation analysis revealed no distinct differences in the interaction between ETS2 and p53-R248Q and that between ETS2 and wild-type p53 (Fig. 3C). Therefore, we next performed an RNA sequencing (RNA-seq) analysis to compare differences in genome-wide gene expression among the p53-wild-type, -KO, -R175H, and -R248Q cells. The RNA-seq data showed that the expression profiles of 572 genes were altered as a result of changes in the p53 status induced by genome-editing (Fig. 3D). Importantly, although the interaction between GOF p53 proteins and ETS2 was markedly different in the p53-R175H and p53-R248Q cells, these cells had similar gene expression profiles. This was confirmed by the principal component analysis (PCA) plot, which showed an overlap between the p53-R175H and p53-R248Q clusters; moreover, these clusters occupied a different region than the clusters of the p53-wild-type and p53-KO cells (Fig. 3E). These data suggest that the function of p53 in the p53-R175H and p53-R248Q mutant isogenic cell lines differed from that in the p53-wild-type and p53-KO cells and behaved as a GOF p53.

### FTD induces aberrant mitotic progression and subsequent apoptosis in tumor cells with *TP53*-GOF mutations

As mentioned earlier, p53 plays a critical role in cell fate decisions in response to FTD-induced DNA replication stress, causing FTD-treated p53-KO cells to undergo apoptosis as a result of aberrant mitosis. To this end, we assessed the response of the p53-R175H and p53-R248Q cell lines (hereafter referred to as p53-GOF cells) to FTD. To visualize the behavior of FTD-treated p53-GOF cells after the G2 phase, we generated p53-GOF cells expressing Fucci SA and performed live-cell imaging. As already reported in our previous study (7), a large proportion of p53-KO cells entered mitosis and exhibited aberrant mitotic progression, whereas the p53-wild-type cells skipped mitosis and transitioned into G1 phase. In p53-GOF cells, the phenotype of mitotic entry and aberrant mitotic progression following FTD treatment was similar to that of p53-KO cells (Fig. 4A, B). Furthermore, the presence of chromosome bridges and the subsequent formation of multi- or micronuclei, which were observed in p53-KO cells after FTD treatment during late mitosis and the next G1 phase, were also highly detected in the p53-GOF cells (Fig. 4C, D, and Supplementary Fig. 3). In addition, nuclear structural defects (i.e., multi- or micronuclei) were also confirmed in the post-aberrant mitotic phase of p53-GOF cells via the immunofluorescent detection of lamina assembly (Fig. 4E, F). FTD-induced apoptosis, as indicated by the accumulation of cleaved PARP and caspase-3, in the p53-GOF and p53-KO cells, but not in p53-wild-type cells because of the transition to senescence (Fig. 4G and (7)). Consistent with this observation, apoptosis was induced in other colon cancer cell lines harboring a different type of p53 missense mutation (Fig. 4H). Collectively, these data indicate that FTD induced defects in mitotic progression (e.g., chromosome bridge formation and cytokinesis failure) and subsequent perturbed nuclear formation and apoptosis, not only in p53-KO cells, but also in p53-GOF cells.

**Figure 4.**
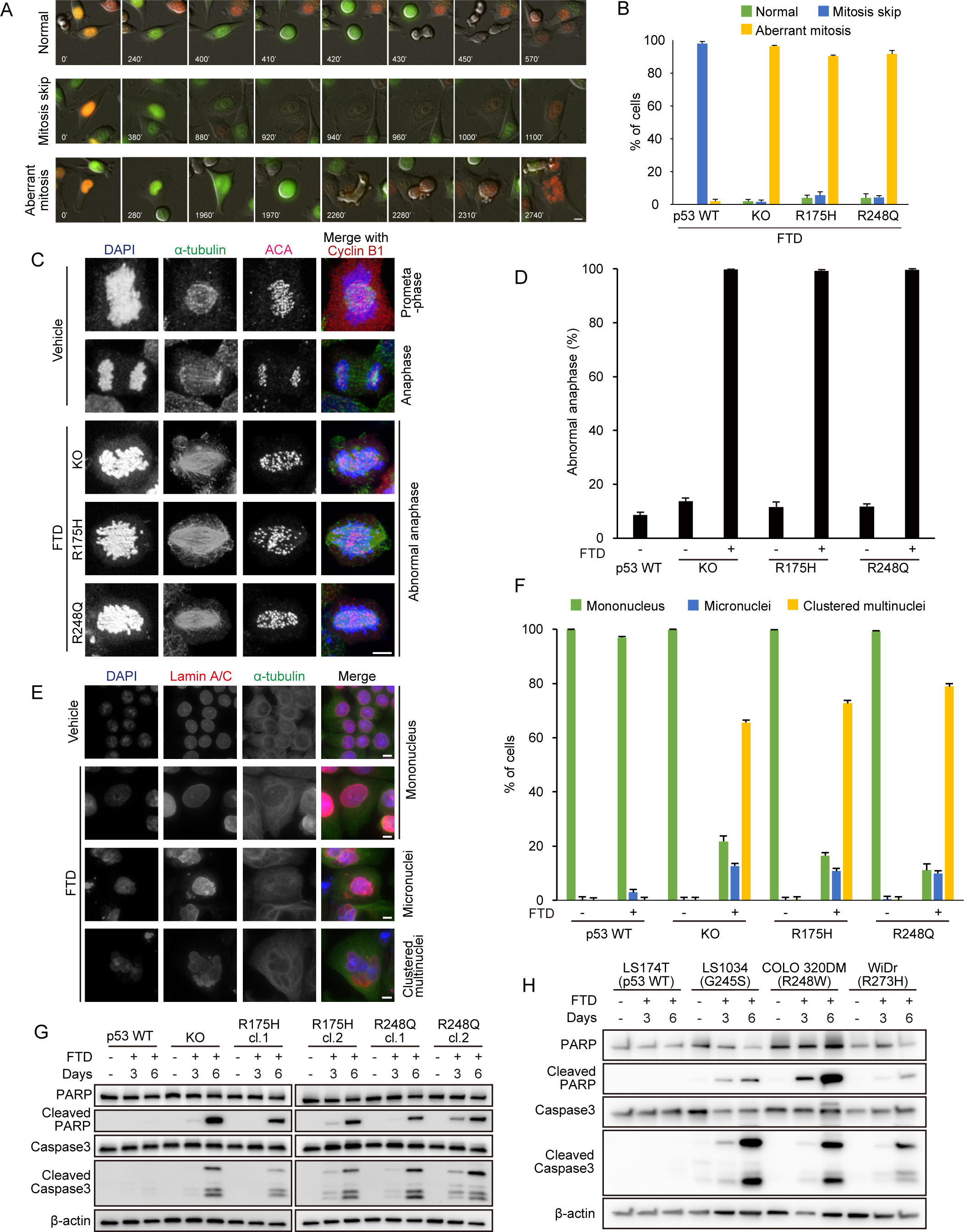
p53 GOF mutant cells display aberrant mitotic progression and apoptotic cell death in the presence of FTD. A and B, Selected frames from live-cell imaging of representative HCT116 mutant cells expressing Fucci SA treated with FTD. Quantification of proportion of cells exhibiting mitosis skip and aberrant mitosis is shown in B. Scale bar, 10 µm. Data are means± s.d. from three independent experiments. C and D, HCT116 p53 mutant cell lines were treated with FTD for 60 hrs, fixed and stained with the indicated antibodies. Representative images of normal or abnormal anaphase are shown in C. Quantification of cells during anaphase with chromosome bridge formation is shown in D. Data are means± s.d. from three independent experiments (≥ 250 cells per experiment). Scale bar, 10 µm. E and F, HCT116 p53 mutant cell lines were treated with FTD for 4 days, fixed and stained with the indicated antibodies. Representative images of nuclei in post-aberrant mitotic phase are shown in E. Quantification of nuclear structure such as multi- or micronuclei formation is shown in F. Data are means± s.d. from three independent experiments (≥ 250 cells per experiment). Scale bar, 10 µm. G and H, FTD-treated cell lines were incubated for the indicated days. HCT116 wild-type and isogenic mutant cells were used in G. Colon cancer cell lines harboring a different type of p53 missense mutation were used in H. Western blot analysis was carried out using antibodies against the indicated proteins.Scale bar, 10 µm.

### FTD exerts an anti-tumor effect on tumor cells bearing *TP53*-GOF missense mutations in tumor models

Finally, we examined how p53-GOF cells responded to FTD and tipiracil (TPI) (FTD/ TPI) treatment in different tumor models. In three-dimensional (3D) cell culture models, FTD treatment suppressed the increase in the size of HCT116 cell spheroids and reduced viability in all cell lines tested (Fig. 5A, B). Furthermore, we assessed the impact of FTD/TPI treatment, which is a clinically approved chemotherapeutic drug combination (3,4), in a xenograft mouse model. We previously reported that FTD/TPI treatment significantly suppressed the growth of HCT116 and HCT116 p53-KO xenograft tumors (7). In the present study, we found that FTD/TPI treatment had a similar effect on p53-GOF xenograft tumors as on those derived from p53-KO cells (Fig. 5C), indicating that FTD/TPI exerted a tumor-suppressive effect irrespective of p53 status.

**Figure 5.**
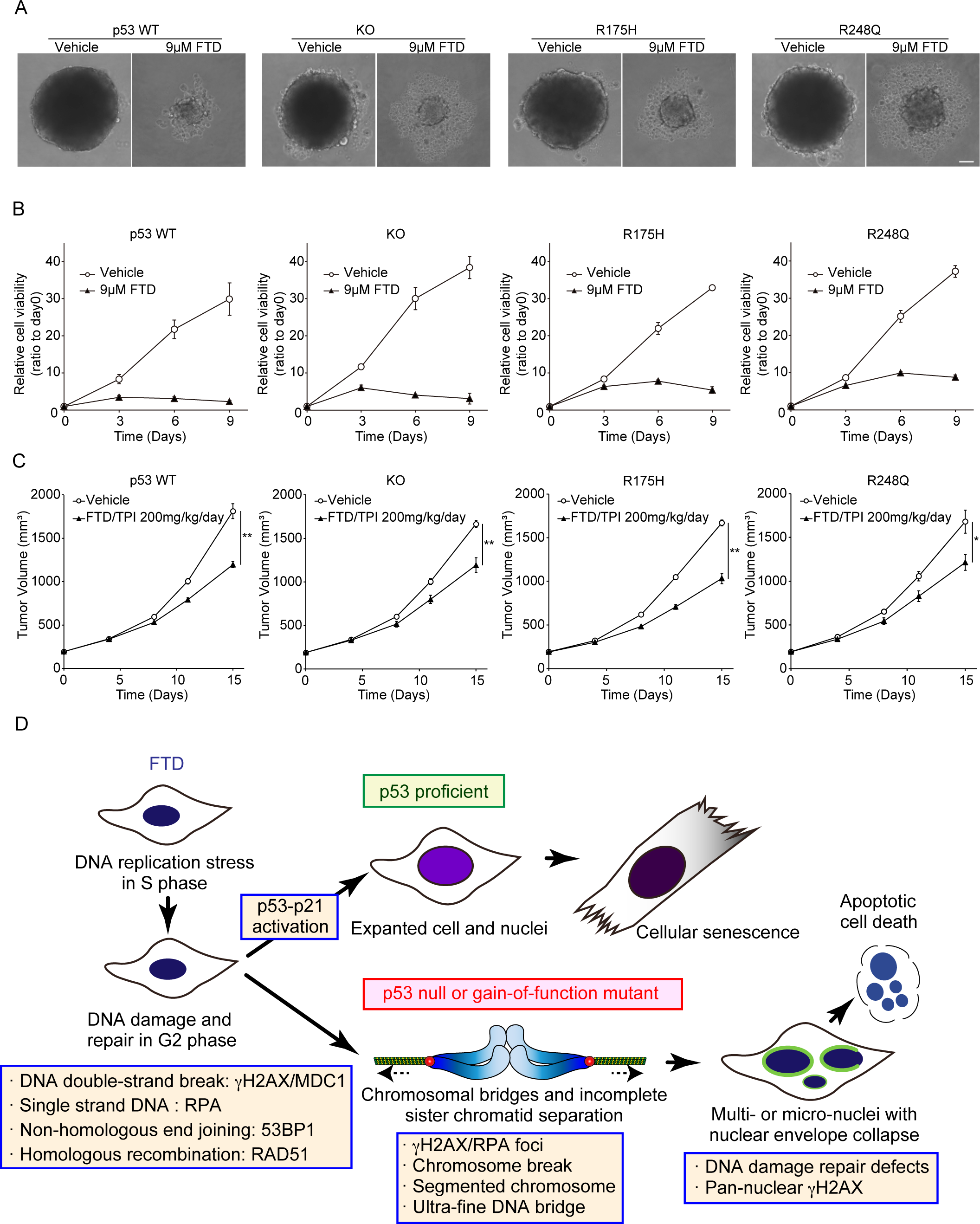
Anti-tumor effect of FTD to cells with different types of p53 status in 3D culture and xenograft models. A and B, Sphere-formation assay using HCT116 cells with indicated p53 status. Representative images of 3D culturing of spheroids in medium with or without FTD are shown in A. Scale bar, 10 µm. Cell viability was analyzed at the indicated times in B. Data are means ± s.d. from three independent experiments. C, Xenograft tumors formed by HCT116 cells with indicated p53 status were subjected to daily FTD/TPI from day 1 to 5 and day 8 to 12. Data are means ± s.e.m. of 6 individual mice. Statistical analysis was carried out at day 15. Aspin-Welch’ s t-test. *: p < 0.05, **: p < 0.01. See materials and methods section for details. C, Model for mechanism of FTD-induced defects in mitotic progression and subsequent apoptotic cell death.

## DISCUSSION

We previously demonstrated that the nucleoside analogue FTD induced DNA replication stress by being incorporated into DNA strands and attenuating replication fork progression; this led to the formation of chromosome bridges during mitosis in p53-KO cells, and ultimately, apoptosis (7,35). In the present study, we identified the underlying mechanisms (Fig. 5D). We found that the FTD-induced DNA damage and unresolved HR intermediates were maintained throughout the G2 phase, mitosis, and the next G1 phase. Moreover, we showed that FTD induced cytotoxicity not only in p53-KO cells, but also in p53-GOF mutant cells. Our findings suggest that FTD treatment induced an insufficient level of DNA replication stress for arrest in the S or G2 phases, which was followed by chromosome mis-segregation during mitosis. Thus, FTD could behave as an effective anti-tumor agent.

We previously performed a biochemical analysis of the *in vitro*-reconstituted DNA replication process, which showed that, at the very least, the replicative polymerases Pol8 and Pol, were implicated in the anti-tumor mechanism of FTD (7). Specifically, we demonstrated that the transient slowing of replication fork progression was caused by FTD incorporation into nascent DNA strands at adenine-rich sites or during replication if a template DNA strand containing tandem repeats of FTD was used as a template (7). Furthermore, FTD-treated cells accumulated RPA-coated ssDNA and contained FANCD2 foci, suggesting that replication fork uncoupling was induced in close proximity to sites of FTD incorporation. Therefore, FTD might induce DNA replication stress responses such as translesion synthesis, template switching, fork reversal, and repriming (36). In accordance, we found that the duration of the S–G2 phase was markedly extended in FTD-treated cells, which was accompanied by RPA-coated ssDNA accumulation during G2 phase (Fig. 1A, 2A, and (7)). Furthermore, the formation of nuclear γH2AX foci was concurrent with RPA-coated ssDNA accumulation. Thus, it is possible that FTD-induced fork uncoupling and subsequent persistent ssDNA accumulation were converted into irreparable DSBs via replication fork collapse (37), the nuclease-mediated cleavage of replication forks (38,39), RPA exhaustion, and replication catastrophe (40). DNA replication stress, induced by aphidicolin, causes the appearance of cytogenetic lesions at common fragile sites (41). Meanwhile, FTD may exert genome-wide effects by inducing DNA replication stress at adenine-rich sites. Although both aphidicolin and FTD affect the rate of replication fork progression and induce DNA replication stress, FTD may do so more efficiently and for longer periods than aphidicolin. This difference may give FTD an advantage over aphidicolin in the process of RAD51-mediated recombination intermediate formation during template switching, fork reversal, and HR (36). Here, we found that the formation of RAD51 foci occurred in G2 phase (Fig. 2A). Furthermore, a large proportion of cells exhibited severe chromosome abnormalities in the subsequent mitosis (Fig. 2F, G). FTD treatment induces chromosomal elongation and segmentation (via defective Holliday junction resolution), which typically occur in cells lacking the nucleases GEN1 and MUS81 (29,42). Our data suggest that FTD induced HR intermediates during late S phase and G2 phase, resulting in chromosome bridge formation (via unresolved recombination intermediates) during anaphase. Because resolvases, such as GEN1 and MUS81, function normally in the HCT116 cells used in this study, FTD-induced HRs may have instead been generated by interference with polymerase ƞ via a D-loop extension (43). Moreover, the recombination intermediate structures induced by FTD might not be recognized as substrates for a strict checkpoint response, which is required to arrest cells at the S-G2 phase, similar to the HR intermediate structure induced by GEN1- and MUS81-depletion (29,42). Indeed, we found that a large proportion of FTD-treated cells entered mitosis and underwent chromosome segmentation (Fig. 2F, G). Such a FTD-induced chromosomal structure should trigger the generation of not only UFBs but also numerous chromosome bridges (visualized by DAPI staining), the end result of which is cytokinesis failure and the subsequent formation of tetraploid G1 cells (Fig. 1B–C, 2H–I). Meanwhile, a ssDNA structure, possibly derived from HR intermediates, was observed during anaphase. Remarkably, this structure can be sheared by a tension of 1–2 nN (44), which is equivalent to the tensile forces generated by spindle microtubules (45). Although the mitotic duration of chromosome congression and subsequent segregation in FTD-treated cells persisted for several hours (Supplementary Fig. 3 and movie), this may have been sufficient to convert ssDNA into DSBs by the pulling force of spindle microtubules. Therefore, our data suggest that FTD-induced mitotic DNA damage is not only carried over from the G2 phase but also arises during mitosis.

FTD-induced apoptosis occurs at least 3 days after FTD administration (7), when most cells are in G1 phase after aberrant mitotic progression. Having navigated abnormal mitosis, most cells were characterized by multi- and/or micronuclei. Micronuclei can be caused by lagging chromosomes (31). In accordance, we found that the FTD-induced micronuclei were encapsulated in the remnants of the disrupted nuclear envelope, failed to regulate the DNA damage response, and had extensive DNA damage. Hence, we concluded that FTD treatment induced the apoptosis of p53-KO cells via the accumulation of DNA damage. Furthermore, micronucleus-derived cytosolic nucleic acids promote the activation of the cyclic GMP–AMP synthase (cGAS)/stimulator of interferon genes (STING) pathway (46). We hypothesized that since FTD treatment induced the formation of micronuclei, it may also activate the cGAS/STING pathway, leading to the increased expression of type I interferon and other inflammatory cytokines. This would ultimately promote the activation and infiltration of anti-tumor immune cells. Indeed, the combination of PARP inhibitor and immune checkpoint inhibitor enhances the anti-tumor response (47). Therefore, an investigation into the potential synergistic effect of FTD and immune checkpoint inhibitors is warranted.

In this study, we delineated the mechanism by which FTD induced cytotoxicity in HCT116 p53-null mutant cells. Because of the high prevalence of *TP53*-targeting missense substitutions in human cancer, we further focused on the behavior of FTD-treated p53 missense mutant cells. We generated genome-edited, p53-missense-mutant, isogenic HCT116 cell lines using the CRISPR/Cas9 system and constructed a p53 mutant cell panel by including p53 wild-type and p53-KO cells. We believe that this model system is a powerful tool for evaluating the effect of p53 status on the cellular response to anti-cancer drugs under conditions that exclude the effects of genetic variations other than p53 status. Importantly, we found that the cytotoxic effect of FTD was similar in all the cell lines tested (Fig. 5A–C), suggesting that FTD exerted its effects in a p53-status-independent manner. Furthermore, this result implies that FTD can overcome the effect of a genome-wide increase in histone methylation and acetylation, caused by the p53-GOF-mediated upregulation of chromatin regulatory genes, including those encoding the methyltransferases MLL1 and MLL2, and the acetyltransferase MOZ (34).

Finally, in this study, we showed that the FTD-induced DNA replication stress and subsequent chromosome bridge formation contributed to cytotoxicity. Chromosome bridges, including UFBs, are a potential source of genome instability. These bridge structures induce the formation of micronuclei, which can trigger a catastrophic genome rearrangement called chromothripsis (48). This phenomenon is related to aggressive tumor behavior (e.g., tumor progression and chemoresistance) and signals a poor prognosis for cancer patients (49,50). However, the FTD-induced chromosome bridges and subsequent micronuclei also trigger apoptosis. These different cell fates seem to be decided by the threshold levels of chromosome breaks and DNA damage, according to the amount of chromosome bridges formed during anaphase. Thus, identifying targets to promote chromosome segregation abnormalities may improve the efficacy of FTD treatment.

## Authors’ Contributions

**T. Wakasa:** Investigation, data curation, formal analysis, validation, visualization. **M. Iimori:** Conceptualization, supervision, investigation, data curation, formal analysis, methodology, project administration, resources, validation, visualization, writing–original draft, writing–review and editing. **K. Nonaka:** Investigation, data curation, validation. **A. Harada:** Methodology, resources. **Y. Ohkawa:** Methodology, resources. **C. Kikutake:** Formal analysis. **M. Suyama:** Formal analysis. **T. Kobunai:** Formal analysis, visualization, validation. **K. Tsunekuni:** Investigation, Validation. **K. Matsuoka:** Investigation, Validation. **Y. Kataoka:** Validation. **H. Ochiiwa:** Validation. **K. Miyadera:** Validation. **T. Sagara:** Validation. **E. Oki:** Validation. **S. Ohdo:** Validation, Supervision. **Y. Maehara:** Supervision, funding acquisition. **H. Kitao:** Supervision, funding acquisition, project administration, writing–review and editing.

## Acknowledgments

We greatly appreciate Dr. Kazumitsu Maehara (Kyushu University) for gene expression analysis, Dr. Sei Shu (Chugai Pharmaceutical Co. Ltd.) for helpful discussions and critical reading, and Dr. Shichao Qiu (Kyushu University) for helpful discussions. We also thank Masako Kosugi, Atsuko Yamaguchi, Naoko Katakura, and Tomomi Takada for their expert technical assistance, and staff at The Research Support Center, Research Center for Human Disease Modelling, Graduate School of Medical Sciences, Kyushu University for technical support. This work was supported by the Ministry of Education, Culture, Sports, Science and Technology of Japan (awarded to Y. Maehara, JSPS KAKENHI grant number 22K19577).

