## Supplemental figure for "The anti-tumor effect of trifluridine via induction of aberrant mitosis is unaffected by mutations modulating p53 activity"

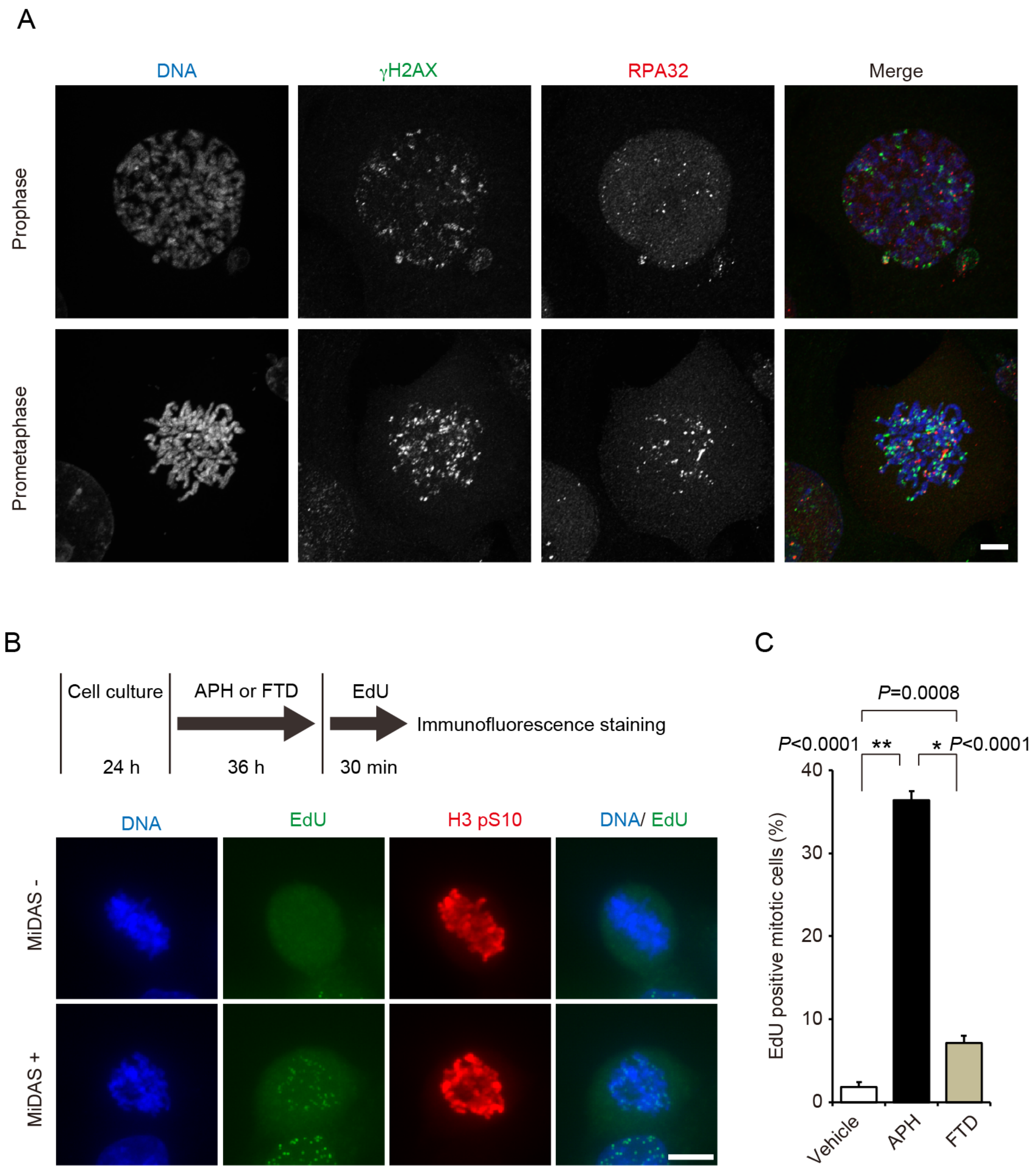


**Supplementary Figure 1. The response of p53-KO cells to FTD treatment during mitosis.**

**A,** Immunofluorescence images showing the localization of gH2AX/ RPA32 in FTD-treated mitotic cells. Cells were fixed and co-stained with the indicated antibodies. Scale bar, 10 µm. **B** and **C,** Representative images of mitotic DNA synthesis (MiDAS) and its experimental scheme are shown in B. Quantification of EdU positive cells is shown in D. APH, aphidicolin. Data are means± s.d. from three independent experiments (≥ 250 cells per experiment). Scale bar, 10µm.

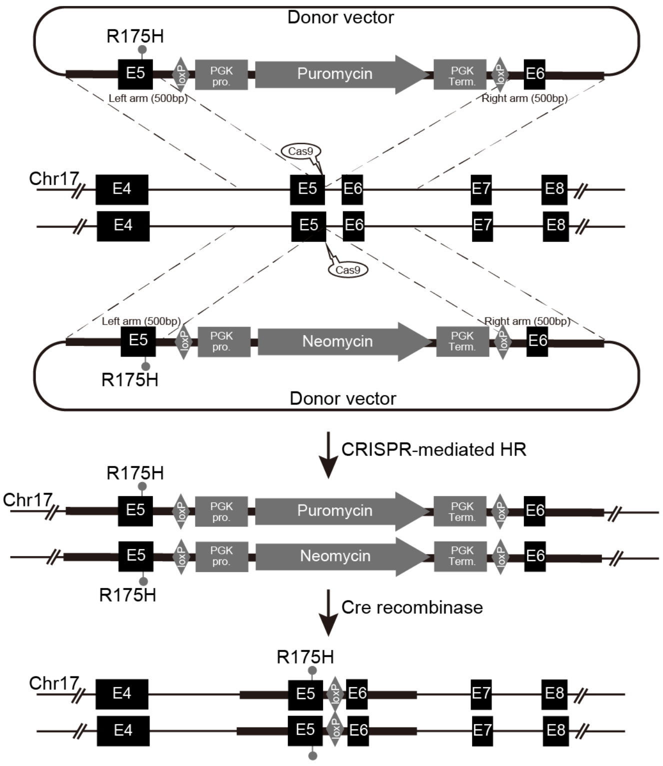


**Supplementary Figure 2. Experimental scheme for generating p53-GOF mutation knock-in cells.**

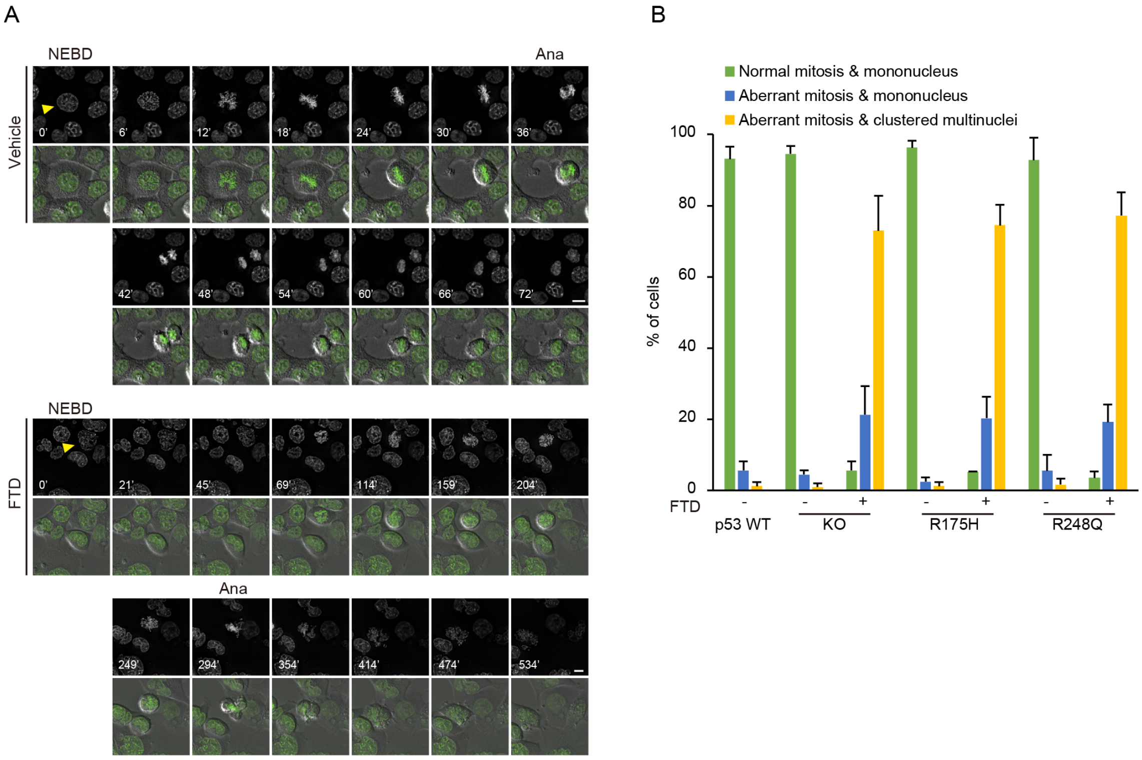


**Supplementary Figure 3. Aberrant mitotic progression following FTD treatment in p53-GOF cells.**

**A,** Selected frames from live-cell imaging of representative HCT116 p53-GOF cells expressing histone H2B-GFP treated with FTD. Time (minutes) after nuclear envelope breakdown (NEBD) is shown on the images. Arrowheads highlight representative cell images. Ana, onset of anaphase. Scale bar, 10 µm. **B,** Quantification of cells exhibiting aberrant mitotic progression and post-mitotic nuclear structure formation. Data are means± s.d. from three independent experiments (≥ 250 cells per experiment). Scale bar, 10µm.
