## Supplemental table for "The anti-tumor effect of trifluridine via induction of aberrant mitosis is unaffected by mutations modulating p53 activity"

Supplemental Table 1. Primer, gRNA and siRNA sequences

| Primer | Sequence (5'-3') | Application |
|---|---|---|
| TP53 R175 sgRNA sense | CACCGGCAGTCACAGCACATGACGG | gRNA for pX330 |
| TP53 R175 sgRNA antisense | AAACCCGTCATGTGCTGTGACTGCC | |
| TP53 R248 sgRNA sense | CACCGGTGTAACAGTTCCTGCATGGG | |
| TP53 R248 sgRNA antisense | AAACCCCATGCAGGAACTGTTACACC | |
| LoxP_PGK pro_F | ATCGATAAGCTTGATGGATCCTAACGCGTCGATCATATTCAATAAC | PCR to construct donor vectors using In-Fusion cloning system |
| LoxP_PGK pro_R | GCTAGCGGTAAGCTTCAGCTGCTCG | |
| PGK ter_LoxP_F | GCTAGCCGCCCGCCCCACGACCCGC | |
| PGK ter_LoxP_R | CTGCAGGAATTCGATGGATCCTAGTGAACCTCTTCGAGGGAC | |
| Neo cassette_F | AAGCTTACCGCTAGCATGATTGAAC | |
| Neo cassette_R | GGGCGGGCGGCTAGCTCAGAAGAAC | |
| Puro cassette_F | AAGCTTACCGCTAGCATGACCGAGT | |
| Puro cassette_R | GGGCGGGCGGCTAGCTCAGGCACCG | |
| R175 up S_F | GCAGCCCGGGGGATCCCTTAGGAGGCTGAGGTGGGAAGATC | |
| R175 up S_R | CGACGCGTTAGGATCCCCTGTCGTCCTCCAGCCCCAGCT | |
| R175 down S_F | AGGTTCACTAGGATCCTGGTTGCCCAGGGTCCCCAGGCCT | |
| R175 down S_R | TAGAACTAGTGGATCCCTAATTTTITITTGTATTTTTCAGT | |
| R248 up S_F | GCAGCCCGGGGGATCCAGTGGCTCATGCCGIAATCCCAGC | |
| R248 up S_R | CGACGCGTTAGGATCCCAGTGGCAGGGGGCAAGTGGCT | |
| R248 down S_F | AGGTICACTAGGATCCCTGCTGTGCCCCAGCCCTGCTIG | |
| R248 down S_R | TAGAACTAGTGGATCCGCATAACTGCACCCTIGGCCCC | |
| TP53_F | TACCAGGGCAGCTACGGTTT | Site-specific Mutagenesis by Overlap Extension |
| TP53_R | ACTGGGGAGGCAGAGTTAGG | |
| p53 R175H_F | GGAGGTTGTGAGGCACTGCCCCCACCATG | |
| p53 R175H_R | CATGGTGGGGGCAGTGCCTCACAACCTCC | |
| p53 R248Q_F | GGGCGGCATGAACCAGAGGCCCATCCTCA | |
| p53 R248Q_R | TGAGGATGGGCCTCTGGTTCATGCCGCCC | |
| QC175_F | CATGGCCATCTACAAGCAAAGTCAACATATGACAGAAGTTGTGAGGCACT | QuickChange Lightning Site-Directed Mutagenesis Kit |
| QC175_R | AGTGCCTCACAACTTCTGTCATATGTTGACTTTGCTTGTAGATGGCCATG | |
| QC248_F | CACTACAACTACATGTGCAATTCAAGTTGTATGGGCGGCATGAACCAGAG | |
| QC248_R | CTCTGGTTCATGCCGCCCATACAACTTGAATTGCACATGTAGTTGTAGTG | |
| p21_F | AGCGATGGAACTTCGACTTTG | Quantitative RT-PCR |
| p21_R | CGAAGTCACCCTCCAGTGGT | |
| MDM2_F | AAATGAATCCCCCCCTTCC | |
| MDM2_R | CACGAAGGGCCCAACATCT | |
| β-actin_F | CTGGCACCACACCTTCTACAATG | |
| β-actin_R | GGCGTACAGGGATAGCACAGC | |

Supplemental Table 2. The list of antibodies used in this study

| Experiment | Antigen | Supplier | Cat # | Dilution |
|---|---|---|---|---|
| Western blot | p53 | abcam | ab32389 | 1:1,000 |
| | p21 | Cell Signaling Technology | 2947 | 1:10,000 |
| | DDDDK(FLAG) | MBL | PM020 | 1:1,000 |
| | PARP | Cell Signaling Technology | 9542 | 1:1,000 |
| | Cleaved PARP | Cell Signaling Technology | 5625 | 1:1,000 |
| | Caspase 3 | Cell Signaling Technology | 9508 | 1:1,000 |
| | Cleaved Caspase 3 | Cell Signaling Technology | 9664 | 1:1,000 |
| | b-actin | Sigma | A5316 | 1:5,000 |
| Immunofluorescence | Pericentrin | abcam | ab4448 | 1:500 |
| | γH2AX | abcam | ab81299 | 1:200 |
| | RPA32 | abcam | ab2175 | 1:200 |
| | 53BP1 | BD Bioscience | 612523 | 1:200 |
| | MDC1 | abcam | ab50003 | 1:200 |
| | centromere (ACA) | Immunovision | HCT-0100 | 1:10,000 |
| | Cyclin B1 | Merk Millipore | 05-373 | 1:500 |
| | α-tubulin | Sigma | T6199 | 1:2,000 |
| | Lamin A/C | abcam | ab238303 | 1:1,000 |
| | Lamin B1 | abcam | ab16048 | 1:200 |
